# Species Tree Estimation from Genome-wide Data with Guenomu

**DOI:** 10.1101/023861

**Authors:** Leonardo de Oliveira Martins, David Posada

## Abstract

The history of particular genes and that of the species that carry them can be different due to different reasons. In particular, gene trees and species trees can truly differ due to well-known evolutionary processes like gene duplication and loss, lateral gene transfer or incomplete lineage sorting. Different species tree reconstruction methods have been developed to take this incongruence into account, which can be divided grossly into supertree and supermatrix approaches. Here, we introduce a new Bayesian hierarchical model that we have recently developed and implemented in the program Guenomu, that considers multiple sources of gene tree/species tree disagreement. Guenomu takes as input the posterior distributions of unrooted gene tree topologies for multiple gene families, in order to estimate the posterior distribution of rooted species tree topologies.

## 1. Estimation of species trees

If we define a species as a group of interbreeding individuals, then a species tree will describe the series of reproductive isolation events leading to the extant composition of species. However, this process seldomly will be in perfect agreement with a randomly chosen gene since the evolutionary model of a gene only takes into account point mutations and insertions/deletions. At the species level, on the other hand, we also observe the duplication of whole genes or chromosomes, acquisition of new material by lateral transfer from unrelated species, as well as the loss of genomic regions. Furthermore within a given extant or ancestral population, the segregation pattern of a particular allele does not always match exactly the speciation events due to ancestral polymorphisms or deep coalescences *(1)*, a phenomena called incomplete lineage sorting. All these biological phenomena, gene duplication and loss, lateral gene transfer and incomplete lineage sorting can lead to gene trees in disagreement with the history of the species, even assuming we can reconstruct the gene phylogenies without error.

Many methods have been developed to infer the evolutionary history of the species as a whole. One strategy is to judiciously select a “core” gene set that can be assumed to be a good representation of the phylogeny of the species, and then proceed with standard phylogenetic reconstruction approaches *(2, 3)*. Other attempts include analysing the phylogenetic profiles (presence or absence of gene families) *(4, 5)*, protein domain organization *(6)*, conserved gene pairs *(7, 8)*, and gene order information *(9, 10)*. In general, the most successful species tree method should make most use of the available genetic information, and therefore should incorporate gene tree reconstruction while accounting for the disagreement between the gene and species phylogenetic histories. Such species methods can be broadly divided between the supertree *(11, 12)* and supermatrix *(13, 14)* approaches, which are represented visually in Figure 1. What we will call a gene family are actually the homologous sets – the collection of sequences that can be aligned and safely assumed to have a common ancestor. Supermatrix methods depend on an exact correspondence between sequences from different genes, such that they can be concatenated. Supertree methods also have been historically limited to one individual from each species per gene family, although recent improvements have made it possible to analyse more general data sets, as the distance matrix-based inference under the coalescent *(15)* or the Gene Tree Parsimony reconciliation *(16, 17)*.

**Figure 1:**
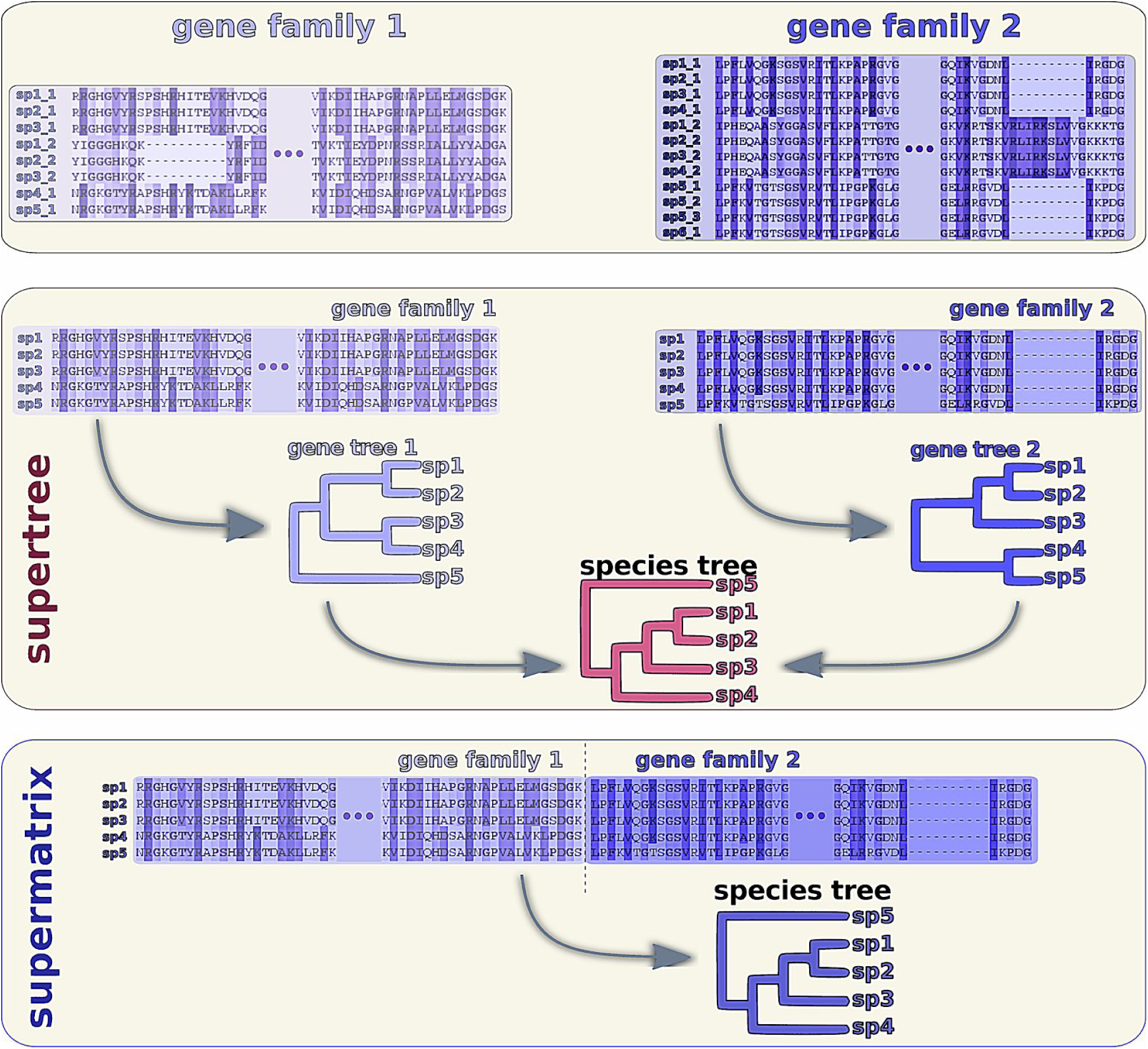
Comparison between the supertree and supermatrix approaches. In the top panel we have an example data set composed of two gene families (homologous sets), already aligned. In this example we have several sequences from the same species in each gene family, which might represent paralogs within an individual sample or more than one sampled individual from the species. A supertree approach (middle panel) would then estimate the gene tree for each alignment independently and then summarise the information from all resulting gene trees into a single tree. A supermatrix approach, on the other hand, would first concatenate all gene family alignments into one single alignment, which would then be used in the phylogenetic reconstruction. Notice how for the supermatrix approach it is essential to find the correspondent sequences for all gene families, a limitation also imposed by several supertree methods. Therefore the user must decide which representative from each species to use, for all gene families. That is, does the sequence *sp1_1* from *gene family 1* corresponds to *sp1_1* or *sp1_2* from *gene family 2*?

## 2. Bayesian inference of species trees

The Bayesian paradigm offers an elegant solution to the problem of reconciling gene trees and species trees, since it makes the hierarchical relation between them explicit. In the same way as the dependence between the gene alignment D and the gene tree G can be be described probabilistically by the phylogenetic likelihood function P(D|G) *(18)*, the gene tree can also be treated as a stochastic parameter depending on a species tree S, as in P(G|S) – where the particular form of P(G|S) varies depending on our assumptions about the G-S disagreement. This will lead to a hierarchical Bayesian model of the form P(S|D) ∝ P(D|G) P(G|S) P(S), where multi-gene data sets can be naturally partitioned as P(S|D) ∝ P(S) ∏ P(Di |Gi) P(Gi |S) (where we exclude other parameters for clarity.)

This idea has been implemented, with more or less success, under a few models – for a review, see *(19)*. A full Bayesian inference model for incomplete lineage sorting has been developed under the multispecies coalescent model *(20, 21)*, which however has a heavy computational burden. Using a birth-death model for the evolution of genes inside the species tree, a Bayesian model for gene tree estimation has been devised *(22)*, being restricted however to a fixed species tree. A maximum-likelihood approach based on a simplified version of this birth-death model has recently been shown to allow for species tree estimation under a hierarchical model *(23)*. Recently, we developed a Bayesian model that takes into account several sources of disagreement between G and S by assuming that P(G|S) follows a multivariate exponential distribution over distances between G and S *(24)*. This distribution works as a penalty against species and gene trees that are very dissimilar under several biologically plausible scenarios. Under this assumption, the model can work with data sets where the orthology relations are unknown, in contrast to the other implementations.

## 3. The software *Guenomu*

Here we will focus on our novel hierarchical Bayesian procedure for species tree estimation, which is part of the Guenomu package (openly available at http://bitbucket.org/leomrtns/guenomu/.) The main program in this package is called `guenomu`, which takes as input a collection of gene trees (posterior distributions or point estimates) and generates a posterior sample of species trees under a multivariate model for the similarity between genes and species trees. At the same time, `guenomu` resamples the gene trees, as well as estimates several other parameters like the distance penalty values and the distance values themselves. The multivariate model assumes that the probability of a species tree generating a given tree is proportional to the similarity between them according to a predefined set of distance measures. Several such distances can be used, and a natural choice are the reconciliation costs: the number of duplications, losses or deep coalescences. That is, the model penalises species trees that must invoke too many duplications or deep coalescences to explain the observed evolution of genes. Another possibility is to use well-known distances even if they lack such a direct biological interpretation, like the Robinson-Foulds distance. In fact `guenomu` can work with all these distances at once, with the only caveat that it neglects branch length information – the multivariate model looks only at the topological disagreement. Another limitation is that it is best suited for distances (or dissimilarity measures, in general) that can work with trees of different sizes, since the gene trees can have several sequences from the same species. To emphasize the generality of these gene data sets, we will call them “gene families” in what follows.

### 3.1 *Guenomu* programs

The Guenomu package is composed of several programs, of which the main one is the Bayesian sampler, called `guenomu`. Besides the main MCMC sampler and analyser, the package also includes a few auxiliary programs that can be executed independently from the main program. We will discuss mainly the Bayesian sampler, but below is a description of other auxiliary functionality offered by the package.

#### 3.1.1 Bayesian sampler

Given a collection of gene families, `guenomu` will estimate the posterior distribution of species trees together with other parameters pertaining to the model. By using a resampling technique, the species tree inference is done through two Bayesian procedures, where first the gene family alignments are used one at a time to sample the distributions of gene trees, and then these individual gene distributions are incorporated into a second, multigene Bayesian model. `guenomu` is responsible for the later, multigene Bayesian sampling, that will sample species trees respecting the distance constraints imposed by the multivariate model. The user is then free to use her favorite single gene phylogenetic inference algorithm to estimate the gene tree distributions (e.g., MrBayes *(25)*, PhyloBayes *(26)*, BEAST *(27)*), that will then be given as input to `guenomu`.

Even gene trees that did not come from a Bayesian analysis can be used in `guenomu`, and a set of compatible species trees will still be inferred. However, besides the probabilistic interpretation being lost in such an attempt, care must be taken to preserve the uncertainty in the gene tree estimation. In other words, even bootstrap replicates should be preferred over point estimates, since `guenomu` relies on the gene tree uncertainty to explore the space of species trees. The primary output of `guenomu` is a file or a set of files in a compact binary format, that must be interpreted by the same program `guenomu` in order to output human-readable files.

#### 3.1.2 Maximum likelihood (ML) estimation of species trees

Usually an MCMC Bayesian sampler can also be used as a ML estimator by simply controlling the temperature of the Monte Carlo chains, as in simulated annealing *(28).* This can be easily achieved by changing a few options in the same program `guenomu`, as we will see shortly.

#### 3.1.3 Summary-statistics coalescent species trees

One important auxiliary functionality of the `Guenomu` package is offered through the program `bmc2_maxtree`, which estimates the species tree under the multispecies coalescent using several distance-based algorithms, namely GLASS *(29)*, SD *(30)*, STEAC *(31)*, and MAC *(15)*. It only needs a file with the species names (as `guenomu`, as we will see below) and the files with the gene trees, and it will output four species tree estimates, one under each modification of the main algorithm. These algorithms rely strongly on the branch lengths of the gene trees, but if they are absent then the program will assume that they are all equal to one.

#### 3.1.4 Pairwise distances between trees

Another useful auxiliary program is `bmc2_tree`, which calculates all pairwise distances between a set of gene trees and a set of species trees. Given two tree files, it will create a file named “pairwise.txt” with a table consisting of the gene tree and species tree indexes, followed by the list of distances between them.

As usual, the program assumes that the names of each species can be found within the gene leaves, and therefore it is sensitive to the order: the gene tree file must be given before the species tree file, in the command line. Furthermore it neglects the root location of the gene trees, since the distances are defined between unrooted gene trees and rooted species trees – hence it does consider the root of the species trees, when defined.

#### 3.1.5 Simulation of tree inference error

The `guenomu` program can benefit from the uncertainty in the estimation of individual gene trees, since it allows the model to consider alternative trees that, however unlikely in isolation, can provide a better fit when analysed in conjunction to other gene families. For each given input gene tree with branch lengths, the auxiliary program `bmc2_addTreeNoise` tries to generate a collection of trees similar to it, based on the premise that shorter branch lengths are more likely to be wrong. In other words, it will transform each tree from an input file into a distribution of trees, increasing the space of possibilities. It also works on trees without branch lengths, and can be used to preprocess the gene tree files estimated by the user before the Bayesian analysis conducted by `guenomu`.

## 4. Input files

The `guenomu` sampler estimates the set of species trees compatible with the phylogenetic information from a set of gene families, which are represented by the so-called input gene tree distributions. These distributions are ideally a good representation of their posterior distributions as estimated by a gene-wise independent Bayesian phylogenetic inference, using state-of-the-art software such as MrBayes *(25)*, PhyloBayes *(26)* or BEAST *(27)*. Therefore previous to the species tree estimation with `guenomu`, the user should already have tree estimates for each gene family.

The idea behind `guenomu` is that the researcher should not have to decide beforehand which genes, partitions or sequences are appropriate – that is, which sequences comprise a locus and which to discard as paralogs, for instance. Therefore, as long as the sequences can be assumed to come from a common ancestor, be it from individuals from a population or from paralogy or speciation, they should be included in the alignment that we call a “gene family”.

`Guenomu` accepts as input gene tree distributions tree files of one of two forms:

1. standard nexus format, where each tree represents a sample from the posterior distribution of a gene-wise independent phylogenetic inference program (like MrBayes, PhyloBayes or BEAST).
1. a compact nexus format, where only topologically distinct trees are represented, together with their representativity (frequency). This file is a valid nexus tree file, but where frequencies (or weights) are annotated as comments alongside each tree. This is e.g. the `.trprobs` file output by MrBayes’ `sumt` command. We will therefore call such files “trprobs” files.

Files from the compact form represent the exact same information as standard nexus files, where the weight values describe how many times each tree was observed in the posterior distribution output by phylogenetic inference programs. Both forms can have the “TRANSLATE” table (from the nexus specification) or not, and branch lengths are allowed despite the fact that they will be ignored by `Guenomu`’s sampler. The trees must not contain multifurcations (except for a deepest trifurcation, characteristic of unrooted trees), and it treats all gene trees as unrooted, even if explicitly defined as rooted.

In practice the input gene tree distributions don’t need to be generated by a Bayesian analysis, and can actually be bootstrap replicates or even a point estimate as the maximum likelihood tree – but we warn that in such cases the probabilistic interpretation is lost.

## 5. Running Guenomu

You can run `guenomu` by including all settings into a control file (and run it like “guenomu run.ctrl”), or by setting the parameters as command line arguments. You can also include the default parameters in the control file, and then overwrite some of these parameters in the command line (which take precedence). The control file can contain a series of arguments, and accepts commentaries inside square brackets, that are then ignored by the program. Absent parameters are replaced by default values.

The only exception is the collection of gene tree files, and a list with the names of all species, that must be given by the user. Many programs demand for a gene-wise mapping between each gene leaf and the species it represents, but `guenomu` only assumes that the species name can be found within the gene leaf name and does the mapping automatically. For instance, if one has for a given gene family, three sequences “a1”, “b1” and “x” all from *Escherichia coli* (they might be paralogous sequences, or individuals from the same locus, or a combination of both), then it is enough to rename the sequences to something like “ecoli_a1”, “ecoli_b1” and “ecoli_x” while including “ecoli” into the list of species. And remembering that in this case all gene families must use the name “ecoli” to refer to this species. It is allowed for a given species not to be found in some gene family, but a gene family member that cannot be mapped to any species in the list will result in error.

Both lists (of gene tree files and species names) can be given as a text file, or added directly into the control file. If included directly in the control file, then they must be between the commands “begin_list_of_” and “end_of_list”, like in the example of Figure 2.

**Figure 2:**
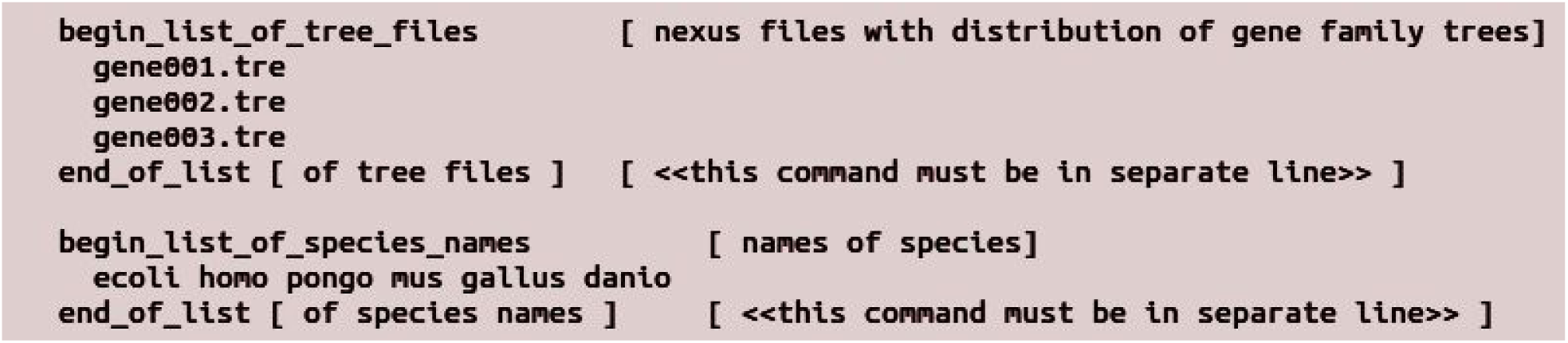
Excerpt of an example control file for `guenomu` showing the list of gene tree files and the list of species names, that should be found within the leaves of the gene trees.

Notice how the comments in brackets are ignored, but still may contain useful tips. Alternatively, if the lists of gene tree files and species names are written in files – let us say, “list_of_trees.txt” and “list_of_species.txt”, respectively –, then these file names can be included in the control file, as shown in Figure 3.

**Figure 3:**
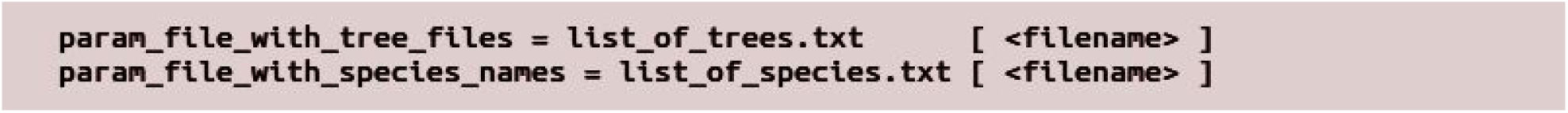
Control file options for `guenomu` that describe the file names containing lists of gene trees and of species names.

Or, equivalently, these file names can be given at run time to `guenomu` as arguments “--genetrees=list_of_genes.txt --species=list_of_species.txt”. With exception of the “begin_list_of_” parameters, all other control file options expect a specific number of values after the equal sign. Other relevant options in the control file are (with examples of values):

- **param_reconciliation_prior = 0.0001**: should be set to a reasonable mean value for the distance penalty parameters. Must be a very small value, to account for our expectation of few disagreements, and furthermore this value is shared across distances (since they are all rescaled to the interval [0,1]). Corresponds exactly to the parameter controlling the hyperhyperprior P(lambda_zero) (see Martins et al., 2014).
- **param_n_generations = 10000 500000**: number of iterations for the burnin stage followed by the number of iterations of the main sampling stage of the Bayesian posterior sampling. In this example, the program will run for 10k iterations without sampling at all, and then it will run for another 500k iterations where now it will sample at regular intervals to compose the posterior distribution.
- **param_n_samples = 1000**: number of posterior samples to save. That is, if we ask for 500k iterations after the burnin (as in the example above), then it will save the state of the chain at every 500 iterations, to form the requested sample size of 1000.
- **param_anneal = 10 1000 0.5 20**: values for the simulated annealing stage that is run even before the burnin stage. They are, respectively, the number of annealing cycles we want, followed by the number of iterations for each cycle, followed by the initial and final (inverse) temperatures of the chain, within each cycle. During this stage no information is saved, unless the program is in “optimization mode”, as we will see later.
- **param_execute_action = importance**: which action should be executed by `guenomu`, to run the importance sampler (regular MCMC run) or to analyse its results, generating human readable output files. The options are then “importance” or “analyse” (“analyze” is also accepted.)
- **param_use_distances = 1111000**: which group of distances should be included in the penalty model, in bit-string representation. The bit-string representation is composed of a one in the n-th position if the n-th distance is used, and zero otherwise. The distances are, from left to right: 1) number of duplications, 2) number of losses, 3) number of deep coalescences, 4) the mulRF distance *(32)*, 5) Hdist_1, 6) Hdist_2, and 7) the approximate SPR distance. The Hdist distances are very experimental, and are based on a matching between branches *(33)*. The approximate SPR distance was developed for detection of recombination *(34)* and may help resolve horizontal gene transfer *(35)*, but is also experimental since, as with the Hdist, it cannot handle the so-called multrees – that is, gene trees with more than one leaf mapping to the same species. That is why in our example these bits are set to zero.

All of the above options have command-line equivalents, which can be consulted by running the program with the help argument “guenomu --help”. For instance, we find that in practice it is easier to set the execution mode for the program at the command-line that at the control file. In this case, instead of setting the param_execute_action option, we run the program with the option “-z 0” for running the MCMC sampler and then run it once more, this time with the option “-z 1” to analyse its output.

### 5.1 Optimization by simulated annealing

If the number of iterations for the main sampling stage of the Bayesian sampling is set to zero, then the program will behave differently: it will assume that we are interested in the simulated annealing results. That is, we are using `guenomu` as an optimization tool, and it will store the chain status at the final iteration of each cycle. The simulated annealing will sample from a ‘modified’ distribution, which is the Bayesian posterior distribution exponentiated to a value (the inverse of its temperature, on a thermodynamic interpretation). In the simulation annealing step, several cycles are performed serially where each cycle is composed of many iterations with an initial and a final temperature. That is, within each cycle the temperature changes across iterations, and the chain state at the end of one cycle is used as the initial state for the next one.

The simulated annealing step was devised to control the initial state of the chain, for the posterior sampling. That is, even within the Bayesian sampling one can benefit from the simulated annealing: if one wants to start the chain at a really random initial value, it is enough to set both the initial and final temperatures to a value below one – which will allow the chain to explore freely even regions of very low probability. This is a safe choice for convergence checks. On the other hand, if one is interested in starting the sample at a good initial estimate of trees, then it might be worth to set the final temperature to a high value, such that only moves increasing the posterior probability are accepted. Increasing the number of cycles allows for the chain to escape local optima.

If one is interested in using `guenomu` as an optimization tool, then we suggest that several cycles of optimization are employed to avoid local optima, and that the initial and final (inverse) temperatures should span a large interval. The output given by `guenomu` (after running it with the option “-z 1”, for instance) will have the optimal estimated species tree at the end of each cycle. We might then look at the most frequent or best species tree overall, remembering that the optimal values are the set of genes and species trees that minimize their overall distances between each other. In such scenario, `guenomu`’s results are therefore a generalization of the SPR supertree approach *(35)*, the mulRF supertree *(32)* or the GTP methods *(17)*, and can be directly comparable by choosing only the appropriate distances.

### 5.2 Output

The main sampler will store its output with all trees and parameters in a binary, compact format that can be not be used by other programs. This file is named “job0.checkpoint.bin”, and if you are running the parallel version then each job will generate its own file, like in “job1.checkpoint.bin”, “job2.checkpoint.bin” etc. Such files must be interpreted by another run of `guenomu` (with option “-z 1”), which will generate a single output file with all numeric parameters as well as the set of posterior species and gene family trees.

#### 5.2.1 Posterior sample of numeric parameters

The output file with the discrete and continuous numeric parameters from all posterior samples is called “params.txt”. One example file is shown in Figure 4 below. It is a tab-formatted file with a header row followed by samples from the posterior distribution, one iteration per row. The header row contains the parameter names, where the order for gene-wise parameters follows the same order as input in the list of gene tree files. Each column represents one variable or statistics (function of the variables) from the posterior sample.

**Figure 4:**
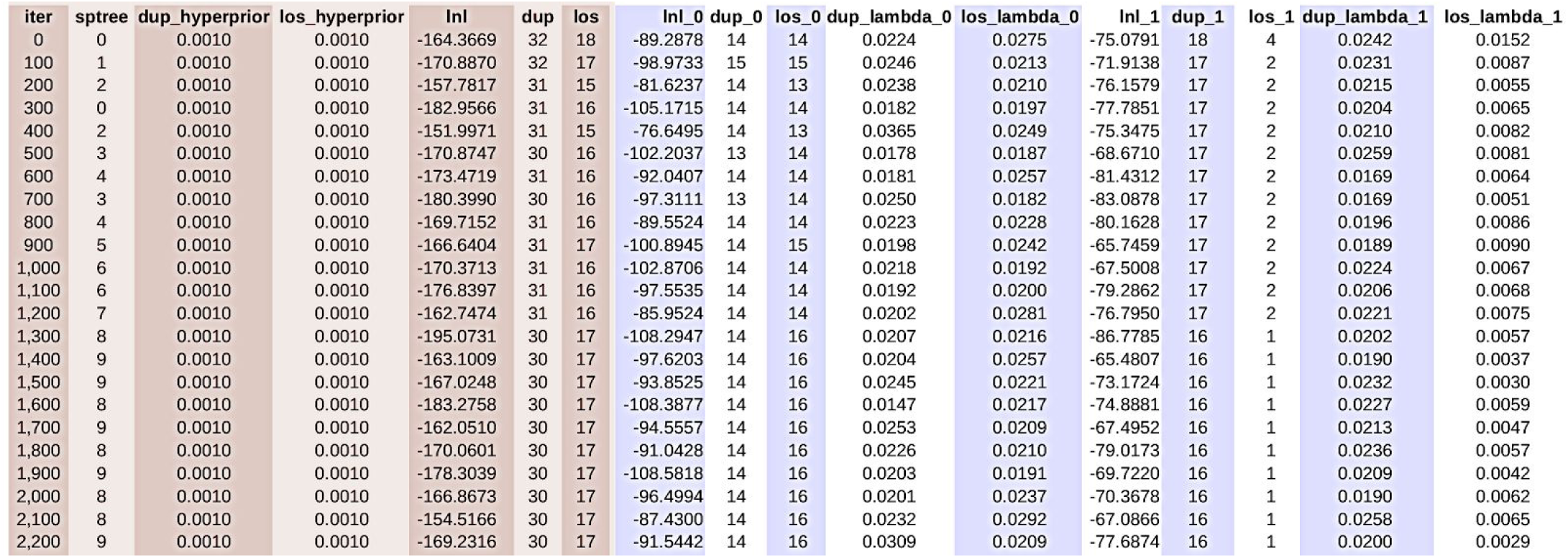
Example of file `params.txt` with posterior samples of parameters from the MCMC chain. The first columns, represented in red, are the global parameters, while the remaining columns represent the parameters per gene family. For this example only duplications and losses were used for two gene families.

Notice that only iterations after the burnin period are sampled, and that furthermore they are already “thinned” to avoid correlation between successive samples – this is accomplished by selecting a number of iterations much larger than the number of posterior samples to save in the parameter input options. The first columns are variables relating to the overall model, followed by gene-wise variables. Therefore the total number of columns depend on the number of gene families being analysed. The overall parameters are, respectively:

1. **iter** – the iteration number of the MCMC run
2. **sptree** – the ID of the species tree, as they appear on output file `species.tre`
3. **dup_hyperprior, los_hyperprior, dco_hyperprior, rfd_hyperprior** – the hyperhyperparameters from the multivariate exponential model, that is, the genome-wise parameters controlling the gene-wise penalties for each distance. The prefixes are the possible distances, namely duplications (“dup”), losses (“los”), deep coalescences (“dco”), and the mulRF distance (“rfd”).
4. **lnl** – the unscaled posterior probability
5. **dup, los, dco, rfd** – the sum of distances over all gene families

While the set of parameters for gene X are:

1. **lnl_X** – the contribution of this gene family to the unscaled posterior probability
2. **dup_X, los_X, dco_X, rfd_X** – the distances between the gene and species trees, for this iteration
3. d**up_lambda_X, los_lambda_X, dco_lambda_X, rfd_lambda_X** – the estimated lambda of the exponential distribution, that is, the penalty for this gene family

This file can be used directly by convergence diagnostic programs like Tracer *(36)* or Coda for R *(37)*, where the behaviour of the global parameters is specially relevant to evaluate if the MCMC has been run for long enough. Within a single run the parameters should appear to be stationary when plotted against the iterations, and their distributions should be similar across independent runs. These are necessary but not sufficient conditions for good convergence, and programs like Tracer and Coda can better assess how the posterior distribution has been explored by our sampler.

#### 5.2.2 Output trees

Besides the output file with numeric parameters, `guenomu` also outputs the resampled gene tree files as well as the posterior distribution of species trees in several formats. The resampled gene trees will contain the same trees as in the input tree files, but with their posterior frequencies – that is, taking into account information from other gene families. These files will have the same name as the input, but with the prefix “post” and in the trprobs format, such that e.g. an input gene tree file named “gene001.tre” will originate a posterior file called “post.gene001.trprobs”.

The posterior species tree distribution is output to three files: “species.tre”, “species.trprobs” and “unrooted.trprobs”. The model in `guenomu` works with rooted species trees, and therefore the files “species.tre” and “species.trprobs” are comprised of rooted species trees. The difference between these two files is the format, where “species.tre” is in standard nexus format and contains all sampled species trees in the order in which they appeared in the MCMC chain. The file “species.trprobs”, however, is in the compact nexus format where only distinct trees are represented, together with their posterior frequencies. The trees inside this file will be named “tree_0”, “tree_1”, etc. where the number is the same as in the “sptree” column of the numeric parameters file, and is used to identify them. This number also appears as comments inside the “species.tre” file, to allow for the mapping of trees between both files. Please be careful since most phylogenetic analysis software cannot handle the trprobs format properly, and therefore the standard nexus file should be used to estimate the consensus tree or in other analyses.

If one is interested in obtaining a consensus tree from the “species.tre” file, then we suggest a few options: 1) the “sumt” option of MrBayes *(25)*; 2) the “consensus()” function of the ape library for R *(38)*; 3) the “consensus” method of the dendropy module for python *(39)*. We do, however, suggest to always check also the posterior frequencies directly from the trprobs files to have an idea about the sharpness of the distributions. On Figure 5 we show an example of the file “species.tre”, that can be compared to the file “species.trprobs” from Figure 6.

**Figure 5:**
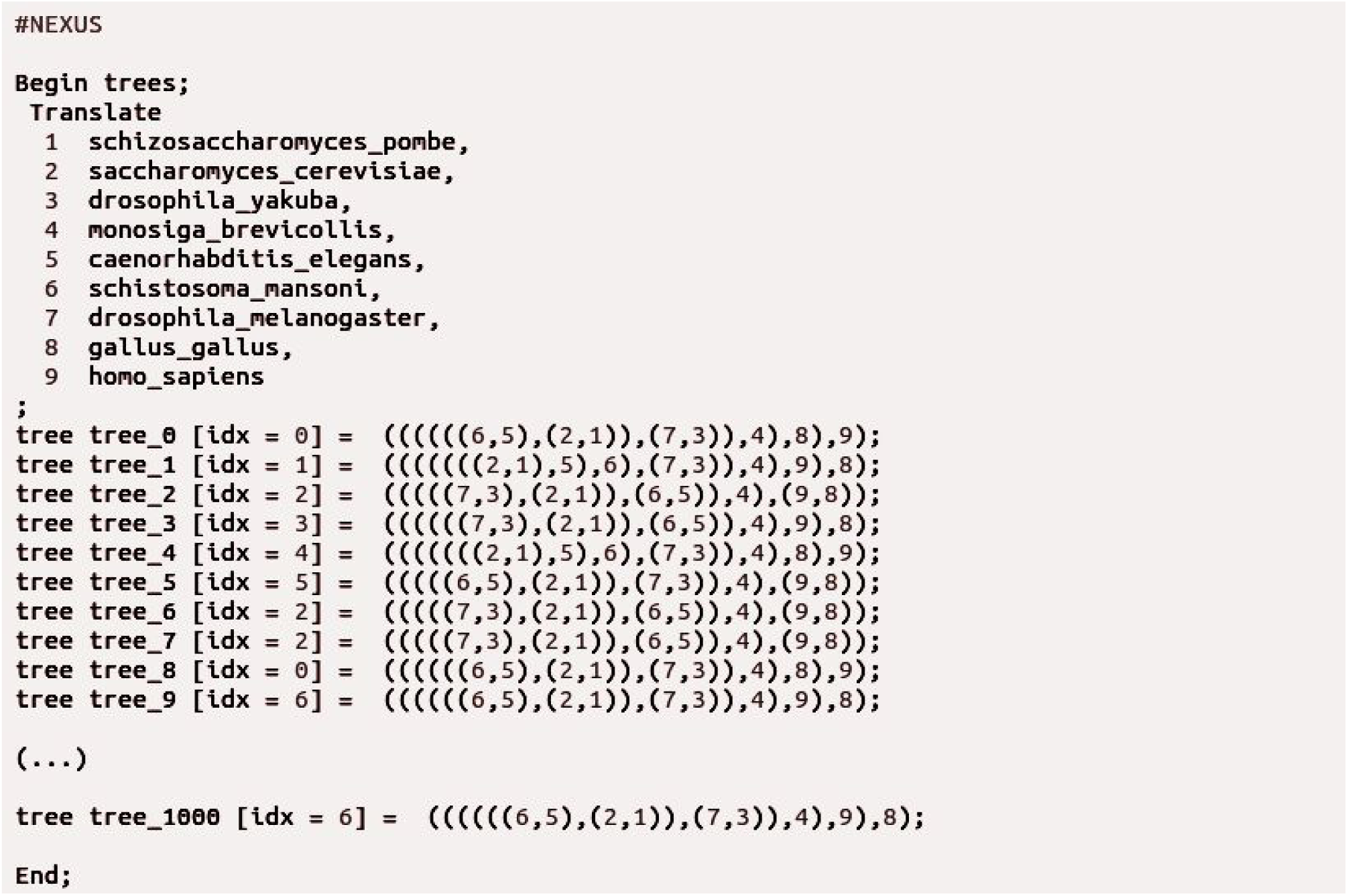
Example of “species.tre” file output by `guenomu`. In this file we have one tree topology per MCMC iteration, where identical trees can be identified by the descriptor “idx” (which is a comment according to the nexus format and therefore does not interfere with other programs.)

**Figure 6:**
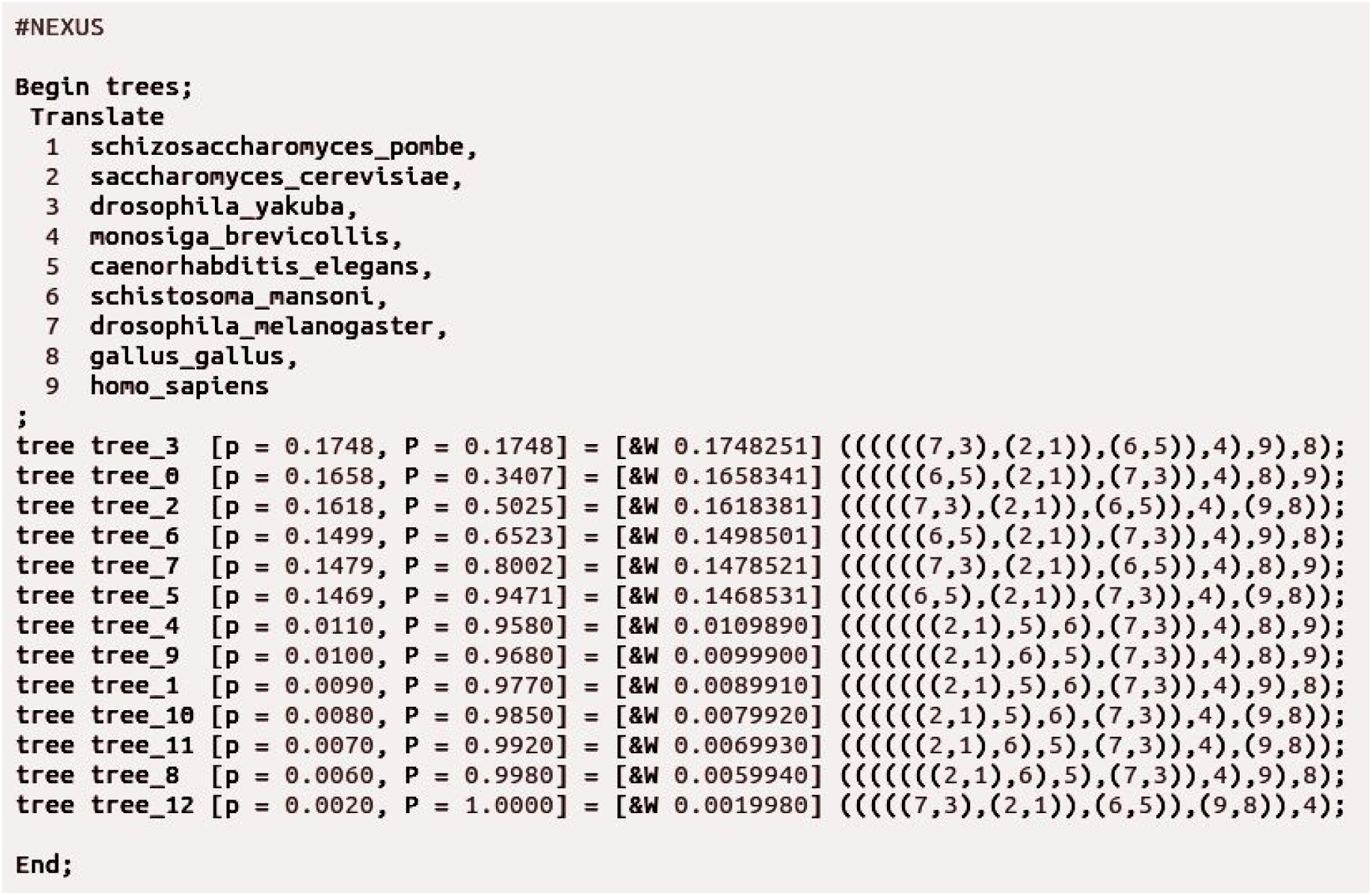
Example of `guenomu`’s “species.trprobs” output file. Unlike the “species.tre” file (see Figure 5), distinct topologies appear only once, and furthermore are ordered according to their posterior frequencies (described within comments.)

Notice how in this file the trees are ordered by frequency, where for instance the tree number 3 is the so-called maximum a posteriori (MAP) tree, with frequency of 17.5%. Notice, also, that some of the trees (e.g. tree_2, tree_3, tree_7 and tree_12) differ only in the root location.

The file “unrooted.trprobs” also contains the posterior distribution of species trees in the compact trprobs format, but this time neglecting the information about the root location. That is, it summarises the posterior distribution assuming that the sampled species trees are unrooted. It is helpful to compare it with “species.trprobs” in order to verify if the samples differ only in their rooting, in which case we might conclude when the data does not allow the model to distinguish between different root locations.In Figure 7 we show the “unrooted.trprobs” file equivalent to the rooted file described in Figure 6.

**Figure 7:**
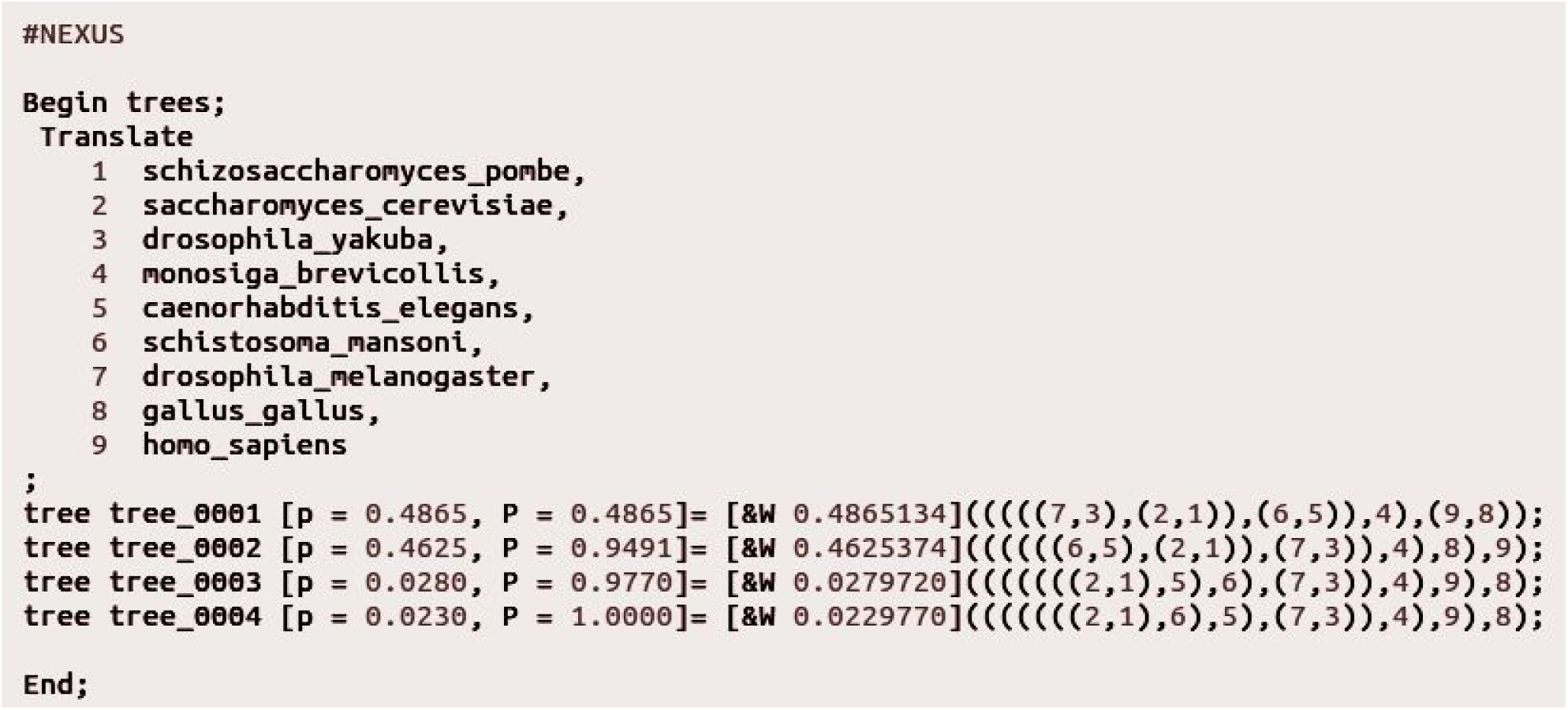
Example of “unrooted.trprobs” output file from the `guenomu` software. This file has the same information as “species.trprobs” (Figure 6) but where the root location of each topology was neglected – therefore joining several trees that differ only in the root location. Notice how the trees are still represented as rooted (deepest node is a dichotomy), just to ease reading and visualization.

The trees are still rooted according to the newick format, but its root node can be safely ignored. In the above example we can see how the information from trees 2, 3, 7 and 12 from file “species.trprobs”all are condensed into the same unrooted tree (tree_0001), with an accumulated frequency of 48.6%

Note that `guenomu` can very fast, such that a single run on a data set with 447 gene family tree distributions and 37 species *(40)* would take less than 6 hours using a single processor.

## 6. Conclusion

The evolutionary history of the species can not be reduced to the phylogeny of a few genes without too many arbitrary assumptions, while the accumulation of data poses new challenges to extracting most information from genomes. Recent Bayesian methods promise to bridge this gap between single-gene and genomic tree inference, while at a high computational cost. On the other hand, novel supertree methods can tackle large data sets, offering however a limited view of the possible evolutionary pathways. Here we show a model, implemented in the program `guenomu`, that tries to integrate strengths from both approaches, allowing at the same time for full utilization of complex data sets under minimal assumptions.

